# Common synaptic inputs are not distributed homogeneously among the motor neurons that innervate synergistic muscles

**DOI:** 10.1101/2022.01.23.477379

**Authors:** A. Del Vecchio, C. Germer, T. M. Kinfe, S. Nuccio, F. Hug, B. Eskofier, D. Farina, R. M. Enoka

## Abstract

The force generated by the muscles involved in an action is produced by common synaptic inputs received by the engaged motor neurons. The purpose of our study was to identify the low-dimensional latent components, defined hereafter as *neural modules*, underlying the discharge rates of the motor units from two knee extensors (vastus medialis and lateralis) and two hand muscles (index and thumb muscles) during isometric contractions. The neural modules were extracted by factor analysis from the pooled motor units and no assumptions were made regarding the orthogonality of the modules or the association between the modules and each muscle. Factor analysis identified two independent neural modules that captured most of the covariance in the discharge rates of the motor units in the synergistic muscles. Although the neural modules were strongly correlated with the discharge rates of motor units in each of the synergistic pair of muscles, not all motor units in a muscle were correlated with the neural module for that muscle. The distribution of motor units across the pair of neural modules differed for each muscle: 80% of the motor units in first dorsal interosseous were more strongly correlated with the neural module for that muscle, whereas the proportion was 70%, 60%, and 45% for the thenar, vastus medialis, and vastus lateralis muscles. All other motor units either belonged to both modules or to the module for the other muscle (15% for vastus lateralis). Based on a simulation of 480 integrate-and-fire neurons receiving independent and common inputs, we demonstrate that factor analysis identifies the three neural modules with high levels of accuracy. Our results indicate that the correlated discharge rates of motor units arise from at least two sources of common synaptic input that are not distributed homogeneously among the motor neurons innervating synergistic muscles.

## Introduction

The motor unit is the final common pathway by which an activation signal is transmitted to muscle and transformed into contractile activity (1). As such, all voluntary actions are accomplished by varying the amount of motor unit activity. Despite early claims to the contrary (2, 3), it is not possible to control the activation of individual motor units (4). Instead, synaptic inputs are distributed broadly among the neurons that comprise a motor nucleus and the motor units that are activated in response to these inputs depends on their relative excitability (5–7). As a consequence of this scheme, the order in which motor units are recruited during a voluntary action is relatively fixed (8–10).

It is the shared synaptic inputs received by the motor neurons that innervate a muscle and not the activity of individual motor units that is responsible for the force it generates (11, 12). In general, the shared inputs can arise from three sources (cortical, brain stem, spinal, and afferent pathways) with varying distributions across different motor nuclei (5, 13–15). One advantage of this scheme is that the shared inputs can engage the motor nuclei of the various muscles involved in an action and thereby facilitate control of the net muscle torque.

It has been hypothesised that the control of multiple muscles is achieved by the activation of sets of motor neurons, that have been referred to as “*neural modules*” or “*motor primitives*” (16–20). Neural modules emerge from common synaptic inputs, or “neural manifolds” (21), that synergistically activate a group of muscles to perform a specific action. For example, evidence from animal studies indicates that the electrical stimulation of spinal interneurons produces coordinated movements that depend on the location of stimulation (22–24). The modularity of neural control in humans has been estimated by measuring the covariation in muscle activation patterns (EMG signals). The modules extracted by factorization analysis have been termed muscle synergies (19, 25) and are assumed to emerge from synaptic inputs that are common to the motor neurons involved in the action.

If the synaptic input is shared among the motor neurons that innervate synergistic muscles, it should generate at least one latent neural module based on the covariance in the discharge times of the activated motor units (21). Previous work has addressed this issue by factorizing EMG signals from different muscles (19, 20, 26–29), which assumes that the motor neurons innervating the synergistic muscles receive similar proportions of common synaptic input from one or more sources.

The neural modules determining coordinated control of multiple muscles can be investigated by pairwise spike train correlations, an approach that gives access to the full statistical operating principles of a neural network (30–32). The purpose of our study was to identify the low-dimensional latent components, defined hereafter as *neural modules*, underlying the discharge rates of the motor units from two knee extensors (vastus medialis and lateralis) and two hand muscles (index and thumb muscles) during isometric contractions (19–21, 33). We hypothesized that the discharge rates of the motor neurons innervating each muscle would be explained by more than one neural module.

We found that the discharge rates of motor units in individual quadriceps and hand muscles could be characterized by two independent muscle-specific neural modules. The discharge rates of most motor units were associated with the neural module for the muscle in which they resided, but others were correlated with either the neural module for the synergistic muscle or both neural modules. We then simulated the delivery of two independent common synaptic currents into integrate-and-fire motor neurons and to validate our approach to identifying latent components. Our findings provide a greater level of detail about the distribution of common synaptic input within and across the motor nuclei that innervate synergistic muscles.

## Results

### Motor unit neural modules

Our approach involved extending the classic method for muscle synergy analysis (17, 19, 20, 25, 28, 34) to motor unit recordings. Instead of treating muscles as individual elements, the discharge times of motor units from different muscles were grouped together. We used a factor analysis that maximizes the correlation between each motor unit and a set of unknown factors, referred to as neural modules. We demonstrate that the factor analysis outperforms other factorization approaches (see Methods), such as principal component analysis and non-negative matrix factorization (19, 35–37), in maximizing the correlation between individual motor unit discharge rates and the latent low-dimensional modules.

Theoretical and indirect experimental observations suggest that a common motor command is distributed to sets of motor nuclei (25, 26, 33, 38–41), which results in the discharge rates of motor units across muscles being strongly correlated during voluntary contractions in humans (42–44) and non-human primates (45). In our approach, we did not assume that there is only one latent signal for each muscle (37, 46, 47). Instead, we decoded populations of motor units from surface EMG signals into series of motor unit discharge times during two tasks that involved the synergistic activation of pairs of muscles.

The experimental setup and correlation analysis for the two vastii muscles is shown in Figure 1. The motor unit discharge times were decomposed with a blind source separation procedure, which identifies each event with no a-priori knowledge on the physiological information conveyed by the individual motor units (48–50). We identified on average across participants 6.9 ± 4.3 and 4.37 ± 2.34 motor units for VL and VM, respectively during isometric contractions at 10% of maximum. As described in the methods, we subsequently smoothed the motor unit discharge times with a Hann window, which retained all the frequencies responsible for muscle force (<5 Hz (11)). As the applied force has a cut-off frequency of ≤20 Hz, the low-pass filtered discharge times (the time series of zeros and ones, Fig. 1C) are strongly correlated with the variance in force during steady contractions (11, 51–53). Consequently, we focused on finding the latent components (i.e., the neural modules) for the low-pass filtered signals (Fig. 1D).

**Figure 1.**
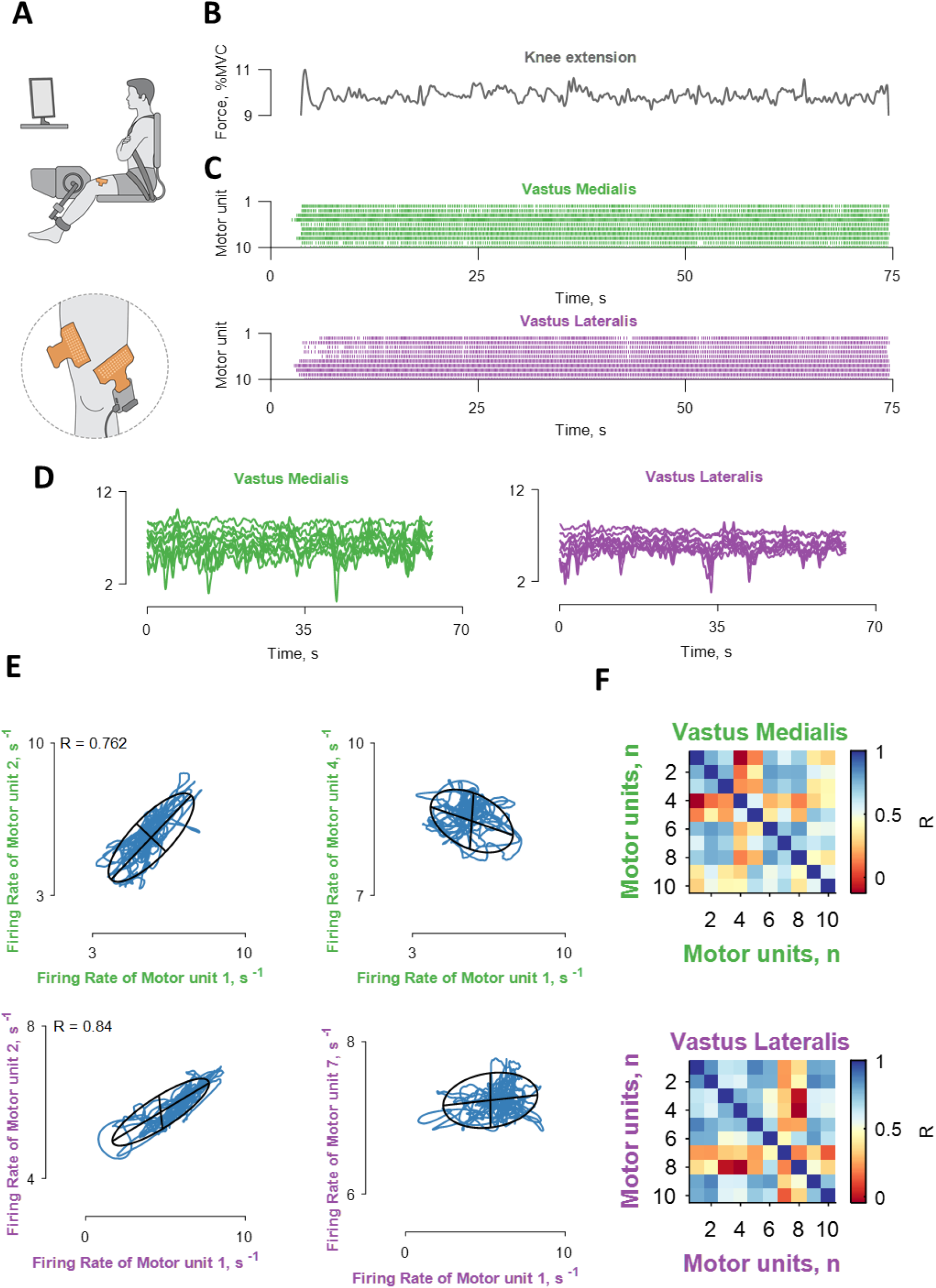
Recordings of muscle force and correlation analysis of motor unit discharge times. **A**. Experimental setup included high-density EMG grids over the vastus lateralis and medialis muscles during isometric contractions at 10% of maximal voluntary contraction (MVC). **B.** The applied force. **C.** The decomposed motor unit discharge times represented in a raster plot for the vastus lateralis (violet) and medialis (green) muscles. **D.** The motor unit discharge times (series of zeros and ones) were convolved with a 400 ms Hann window, which retains the motor unit oscillations responsible for the fluctuations in force during steady contractions. **E.** Four bivariate correlations between different motor units belonging to the same motor nuclei (the labels are color-coded with respect to the muscle as indicated in panels C and D). The blue lines indicate the smoothed discharge rates during the steady-state contraction. **F.** Confusion matrix of the correlation strength between all the identified motor units for the two muscles. Note that the discharge rates within each homonymous motor nucleus exhibited a range of correlation values. R = correlation strength, for both correlations the Pearson’s value was <0.0001.

After converting the discharge times to rates and smoothing the signal, we computed pairwise correlations between each motor unit within the same muscle (Fig. 1E-F). We consistently found correlated and uncorrelated discharge rates of some motor units from the same vastus muscle, which indirectly indicates that not all motor neurons received the same common input (30, 31). Because most previous studies report high correlations among motor units within a motor nucleus (26, 42, 54), it was necessary to assess the level of cross-talk between muscles. Based on a recently developed method (55, 56), we found that the identified motor units had action-potential amplitudes that were statistically similar to the other units in the homonymous muscle and, therefore, were not the result of cross-talk (see Methods).

The average number of identified motor units for the hand muscles was 12.2 ± 3.0 and 4.3 ± 1.2 for the first dorsal interosseous and thenar muscles, respectively, across participants. In contrast to the vastii muscles, Figure 2 shows that the discharge rates of all motor units in each hand muscle were strongly correlated (>0.9), and there were few cases of low correlations (see cluster analysis below). Because of differences in the strength of the correlations between motor units in the vastii and hand muscles, we then examined the latent components between motor units across participants and muscles with a factorization approach.

**Figure 2.**
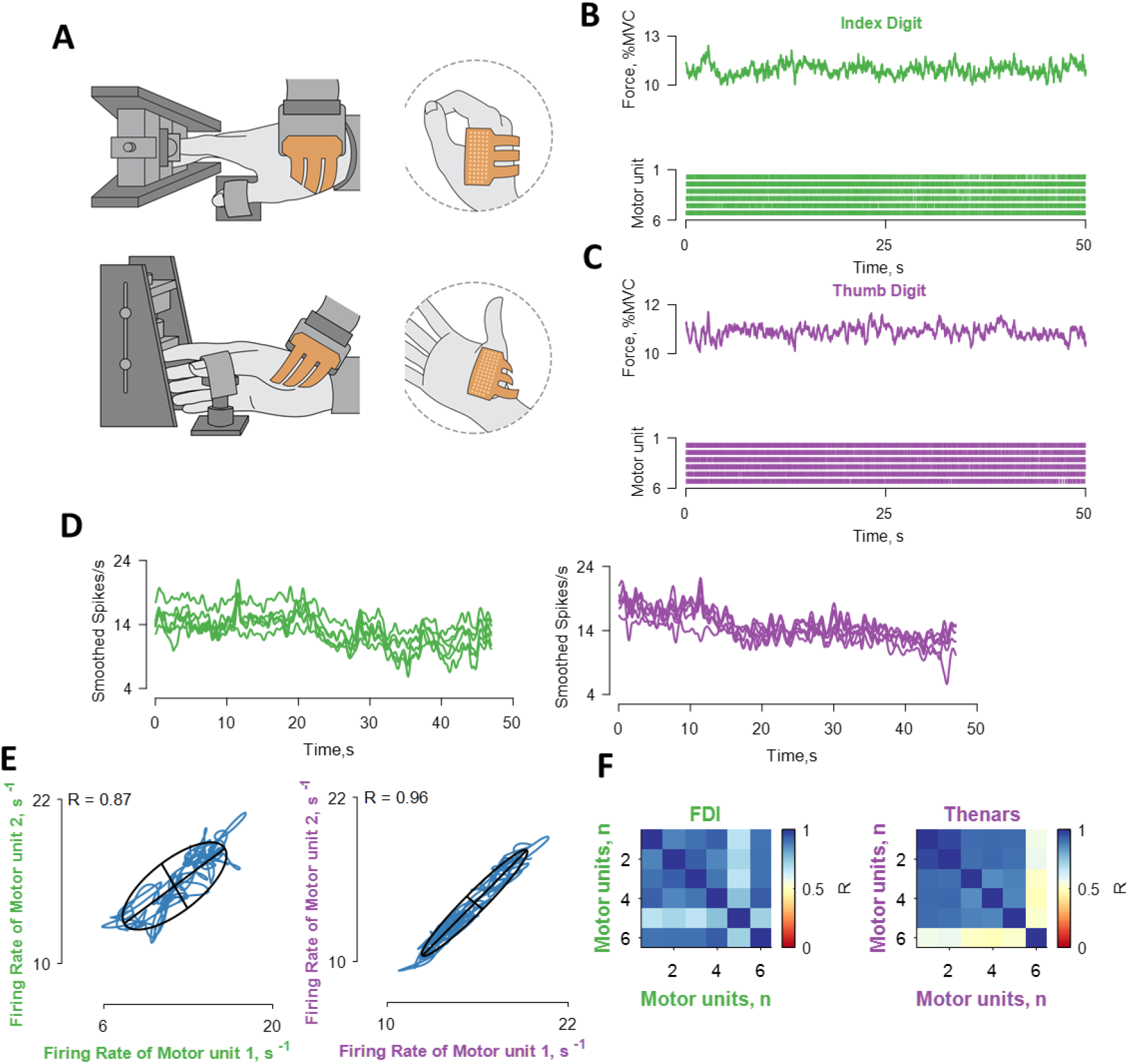
Recordings of muscle force and correlation analysis between the discharge times of motor units in hand muscles. **A.** Experimental setup involved high-density EMG grids placed over the first dorsal interosseous and thenar muscles. **B-C.** The applied force and the discharge times of motor units shown in a raster plot for the first dorsal interosseous (green) and thenar muscles (violet). **D** The motor unit discharge rates (series of zeros and ones) were convolved with a 2.5 s Hann window. **E.** Two bivariate correlations between different motor units belonging to the same motor nucleus (the labels are color-coded with respect to the muscle as indicated in panels C and D). The blue lines indicate the smoothed discharge rates during the steady-state contraction. **F.** Confusion matrix of the correlation strength between all the identified motor units the two muscles. Note that all motor units are highly correlated within each motor nucleus. R = correlation strength, for both correlations the Pearson’s value was <0.0001.

### Factor analysis reveals a distinct organization of common synaptic inputs

Factorization analysis identifies the latent components that covary among sets of variables. This method enables the identification of the potential ‘neural modules’. Figure 3 shows the results obtained from the factor analysis for the vastii muscles of two participants. The factor analysis was applied to all motor units from both muscles; therefore, the extracted neural modules did not have any a-priori muscle-specific constraint. The latent neural modules are superimposed on each muscle (grey lines indicate individual motor unit discharge rates from that muscle). The first two modules explained most of the signal (>80%); therefore, we only used these two factors in the subsequent analysis.

**Figure 3.**
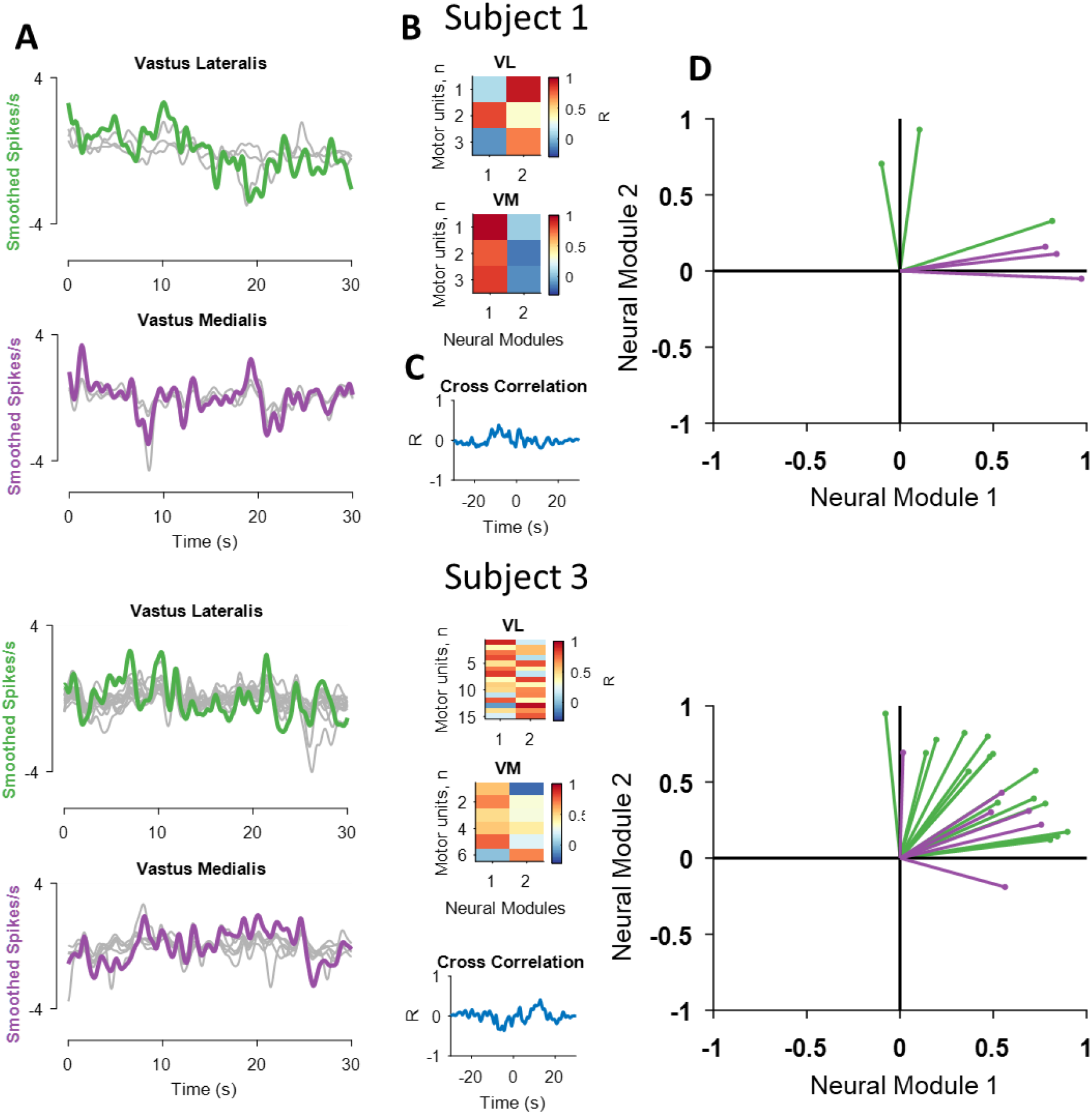
Results of the factorization analysis for the vastii muscles of two subjects. **A.** The smoothed motor unit discharge rates (grey lines, with the mean = 0 spikes/s) and two neural modules derived from the factor analysis (green for the vastus lateralis and violet for vastus medialis). Note the high correlation between the two factors and the discharge rates of some, but not all, motor units for the two subjects. **B.** The two factors were then correlated with the smoothed discharge rates of all motor units. **C.** The cross-correlation values between the two modules. **D.** Projections of the bivariate correlation values for each motor unit with respect to the neural modules. Values close to 1 indicate that a motor unit carries ~100% of the respective module. Note that some vastus lateralis and medialis motor units invade the territory of the other neural module; for example, intermingling of the green and violet lines for Subject 3. Also note that some motor units are only correlated with one module.

We then determined the level of correlation between the discharge rate of each motor unit and the two neural modules (Fig. 3B). This analysis shows, for example, that motor unit number 2 in vastus lateralis for Subject 1 had a stronger correlation with the first neural module, whereas the two other motor units were more correlated with the second factor (Fig. 3D). However, the two modules were not correlated (Fig. 3C). Projections of the two modules (Fig. 3D) indicated that one motor unit in vastus lateralis was located in the module of the vastus medialis motor units. Subject 3 exhibited more intermingling of the motor unit data in the space of the two neural modules (Fig. 3 lower right graph).

We then looked at the overall distribution of the identified motor units within each neural module across participants for the vastii (n=8) and hand muscles (n=8) (Fig. 4). We found that although many motor units from each muscle shared the same module (Fig. 4A-D), the discharge rates of some motor units were correlated with both neural modules. There were also some motor units that showed negative correlations with one of the modules. These negative correlations were more common for the hand muscles.

**Figure 4.**
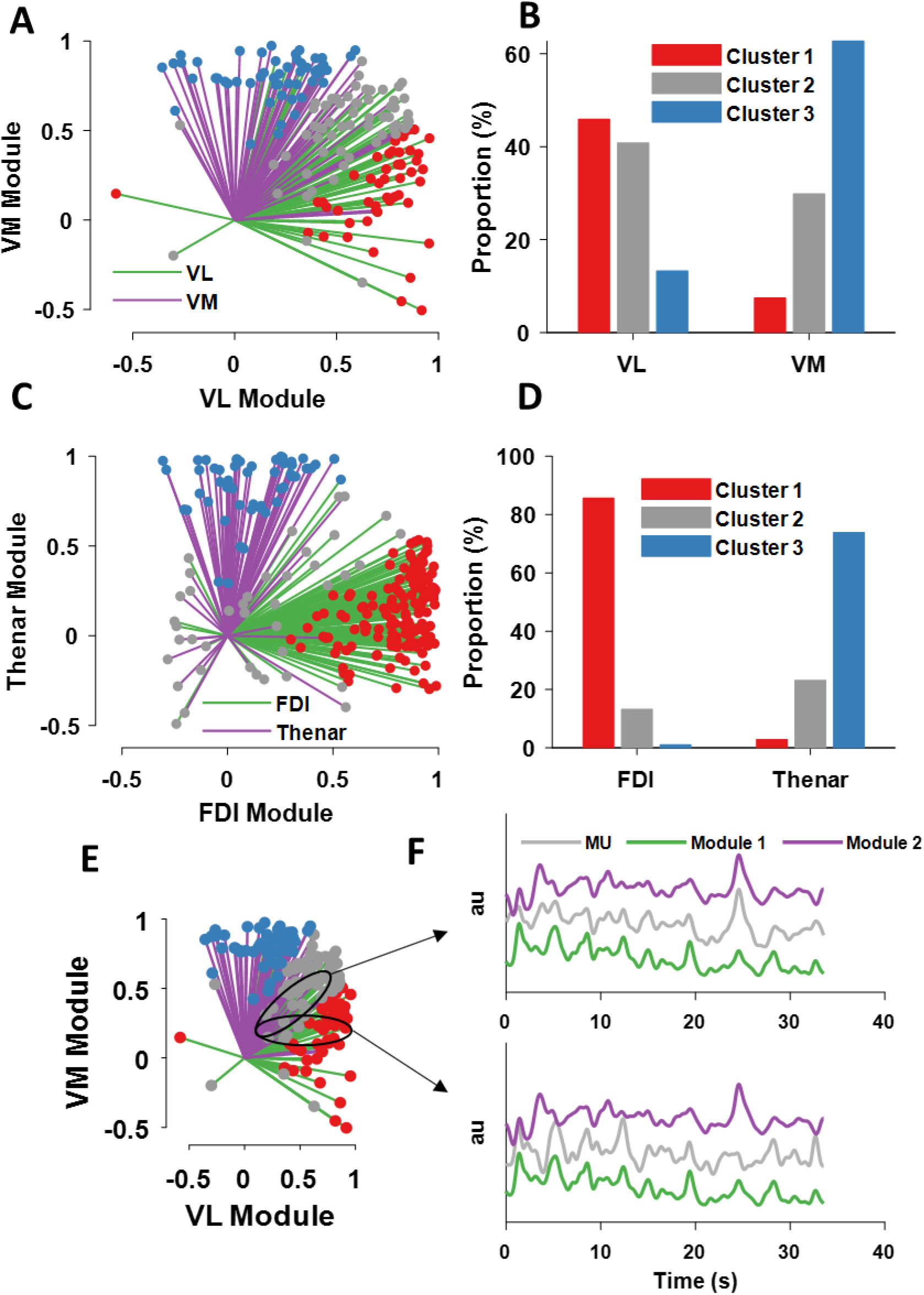
The output of the factor and cluster analysis across all subjects and motor units. **A.** The motor units from vastus lateralis (green) and vastus medialis (violet). Each line indicates the strength (line length) and sign of the correlation between the discharge rate of one motor unit and the neural module. Note that some motor units shared the same module space (indicated in grey dots), whereas others diverged from synergistic control (blue and red) and a few invaded the territory of the other muscle. **B.** A cluster analysis identified three main clusters. Note the grey cluster that indicates the percentage of motor units that shared both neural modules. **C-D**. The same analysis as in A-B but for the hand muscles. Note the smaller proportion of motor units belonging to the shared (grey) cluster in comparison with the vastii muscles. **E-F** An example of two motor units that occupy different module space. The black ellipse is a visual guidance of the territory occupied by the motor unit 1 (top panel in F), which shows the firing rate of that unit correlated to both module 1 and module 2. In contrast, the lower panel in F shows a motor unit that is only correlated with module 1.

We clustered the correlation values of the motor units with the respective modules based on specific centroids (x and y coordinates: [0.66 1.00], [0.40 0.40], [1.00 0.65]), (Fig. 4 B-D). The largest cluster for all muscles was the group belonging to the homonymous muscle; that is, a motor nucleus-specific cluster. Interestingly, the proportion of motor units belonging to the shared cluster was greater for vastus lateralis than vastus medialis (Fig. 4B). The motor nucleus-specific cluster was stronger for the hand muscles, with few motor units present in the shared cluster (<20% for both first dorsal interosseous and thenar muscles). Moreover, there were some motor units for both sets of muscles that diverged from the homonymous control and were more correlated with the other synergistic muscle. This was more evident for the vastii (>10% of motor units) than hand muscles (<3% of motor units).

We then removed the motor units that shared both neural modules and performed coherence analysis between the motor pools. We found approximately a two-fold decrease in the coherence value without the common motor units. For some subjects, the coherence in any of the physiological bandwidths (0-50 Hz), did not differ than for frequencies >50 Hz, which means that there was no coherence between motor units that did not share the same neural module. Conversely, the coherence for the motor units that shared both modules was similar to what previously reported (26, 44). This finding strongly indicates that previous coherence found between muscles from thigh and from the hand is due to the shared inputs that generate the significant coherence value (26, 56).

### Integrate and fire model: factor analysis accurately reflects the interplay of two common inputs

We performed computer simulations to generate a dataset of motor neuron discharge times to determine the optimal convergence and accuracy of factorization analysis on the extraction of neural modules from motor unit data. The aim was to assess the influence of known distributions of two synaptic inputs (*I*_syn1,2_) and their average *I*_syn3_ = (*I*_syn1_ + *I*_syn2_)/2 + independent inputs on the number of identifiable neural modules. The common and independent inputs, as well as the spike times, were approximated by tuning the parameters of an integrate-and-fire model (32, 57).

Because we have no information on the dimensions of the latent components, which reflect common inhibitory and excitatory synaptic inputs, we can model these inputs realistically with an integrate-and-fire model (32, 57) and study the outputs with pairwise correlations and factor analysis. Moreover, the model allowed us to test the influence of time (total duration of spike times), net synaptic currents, and the strengths of the common and independent inputs.

We simulated 480 integrate-and-fire neurons that were activated by applying an independent input and a common input. Two-thirds of the population of neurons received the uncorrelated inputs, *I_syn1_* and *I_syn2_*, and a one-third received *I_syn3_*, which represented the average of the two other inputs (Fig. 5A). We then looked at the correlations between the inputs and outputs (smoothed motor unit discharge rates) as well as the average and standard deviation of the neural modules extracted by the factorization method (Fig. 5). Due to the influence of low-pass filtering of the discharge rates, there was a strong influence of trial duration with the 10-s data being unable to distinguish between the unique and shared inputs. With longer time signals, we were able to retrieve the full dimensions of the three common synaptic input signals by increasing the simulation to 50 s or 80 s (Fig 5. H-I), independently of common and independent input strength.

**Figure 5.**
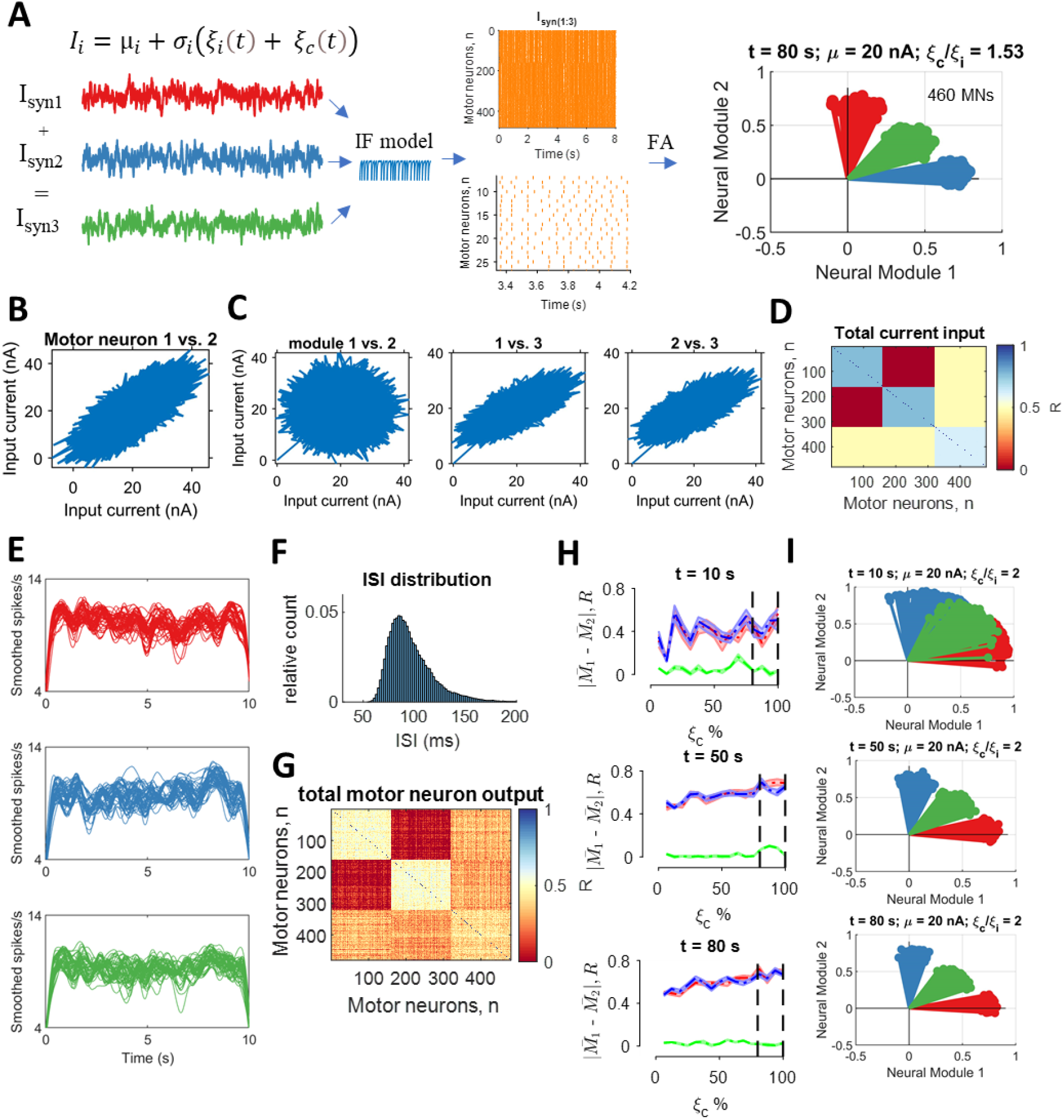
Integrate-and-fire model. We injected correlated and uncorrelated fluctuating currents (*I_i_*) into 480 motor neurons. Two-thirds of the population received two distinct common inputs (*ξ_c_*), and the one-third received the average of the two common inputs plus its own independent noise. The proportion of common and independent inputs ranged in values to reflect the cross-correlation values observed in the experimental data. Similarly, the injected current (20 nA) generated interspike intervals that matched *in-vivo* motor unit data. Each neuron received gaussian synaptic noise reflecting its unique connections (independent input, *ξ_i_*). **A.** μ_*i*_ is the temporal average of the current (20 nA) and *σ_i_* sets the global network state. Raster plot (orange lines) showing some of the data from an 80-s simulation with the proportion of common-to-independent inputs set at 1.53. Note the output of the factorization analysis clearly depicting the space of three injected currents (top right graph), as observed in the experimental recordings. **B.** Pairwise correlation for the first neural module between motor neurons 1 and 2 from a pool of 160 neurons that each received *I_syn1_*. **C.** The averaged total current across all cells plotted between the neural modules. Note that module 1 and module 2 are uncorrelated, whereas there was a high correlation with module 3 due to the shared averaged synaptic current. **D.** Confusion matrix of the correlation strength across all 480 neurons. **E.** The output of the integrate-and-fire model was low-pass filtered at 25 Hz for a 10-s trial. The first and last second was excluded when calculating the correlations to avoid the influence of spike frequency adaptation. Each line corresponds to one motor neuron. **F.** Distribution of interspike intervals across all 480 neurons for an 80-s trial. **H.** Accuracy of factor analysis computed as the average difference of all neurons belonging to each module (|*M_1_ - M_2_*|) at three time points during the simulation. The absolute difference between the modules corresponds to the accuracy of factor analysis in converging in that specific module. The values for the shared module (green) were close to 0, which indicates perfect separation from the two modules. We injected low percentages of common input (0% indicates that the common and independent input are the same). The dashed vertical lines indicate the range of values that reflect in-vivo motor unit correlation values. There was a strong influence of time, so that 10 s of data were insufficient to obtain reliable estimates of the proportion of common input, whereas there were no differences for the data at 50 s and 80 s. **I**. Three representative neural modules extracted by factor analysis at three time points (10, 50, and 80 s) when the common input was twice as much as the independent input.

## Discussion

We analysed the correlation between the discharge times of motor units from different synergistic muscles during isometric contractions with the knee extensors and index finger and thumb muscles. We found two neural modules for the motor units of the vastii muscles, which contrasts with previous findings of only one dominant common input to individual (42) or synergistic muscles (26). As shown in Figure 6, large groups of motor units innervating the VL and VM muscles were associated with specific neural module for each of these muscles, but some motor units were associated with both neural modules. In contrast, fewer motor units innervating the hand muscles (<20%) were associated with both neural modules. Moreover, the discharge rates of some motor units were not correlated with the neural module for the muscle in which they reside, but instead were correlated with the neural module for the synergistic muscle.

**Figure 6.**
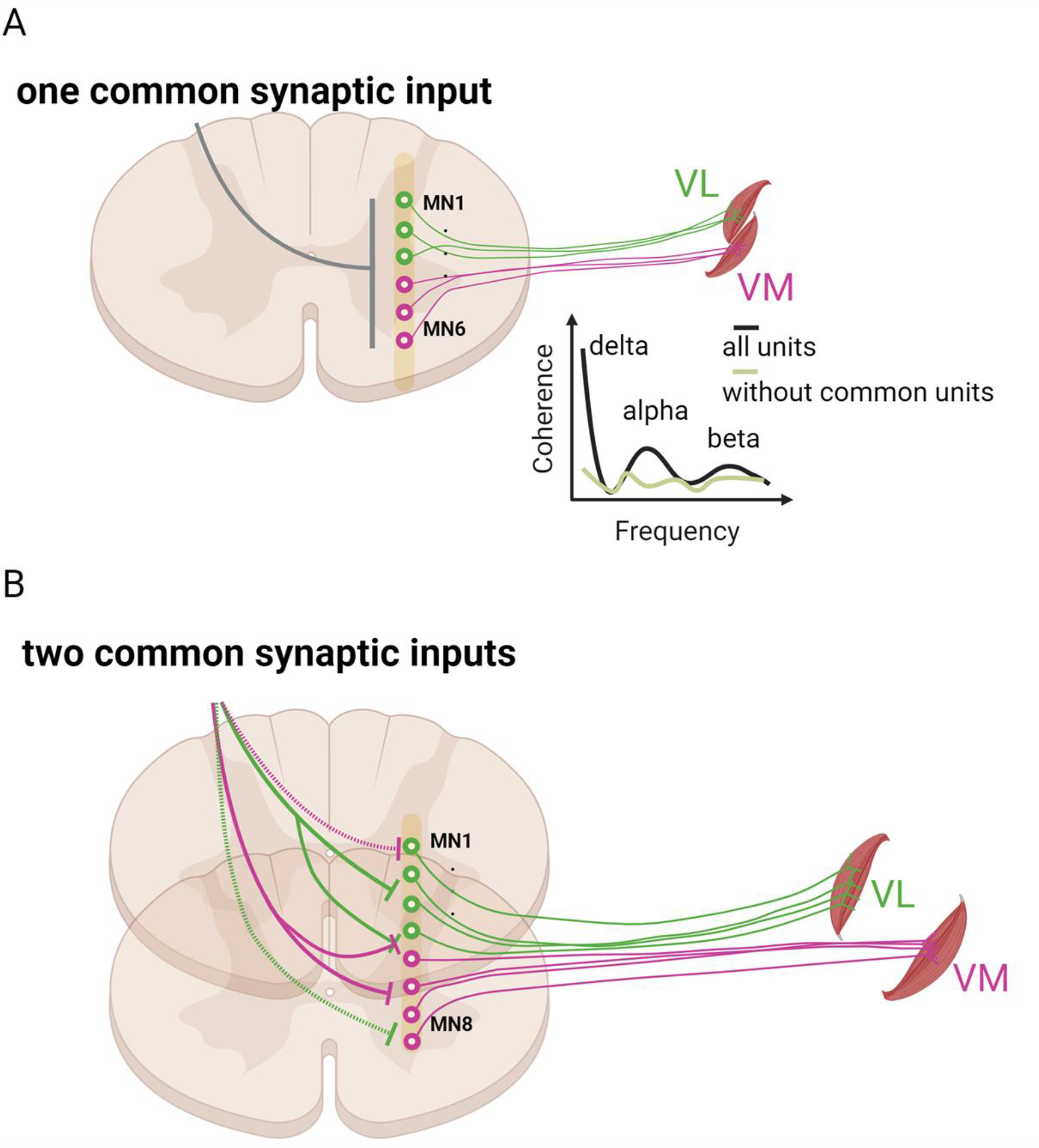
Schematic representation of the results and suggested neural connectivity of voluntary motor commands to motor neuron pools. **A.** Previous studies that have grouped the vastii or hand muscles based on global EMG signals have found strong coherence between the two muscles, indicating a unique common input to the synergistic muscles. After we removed those motor units that shared both neural modules, the correlation between the two pools of motor units was significantly reduced, as indicated in the coherence graph. This indicates that the coherence found in previous studies is mainly attributable to those motor units that shared two distinct sources of common synaptic. **B**. Visual representation of our current findings. With factorization dimensionality techniques, we found that there are at least two sources of common synaptic input to motor neurons that innervate the vastii and hand muscles. Most motor neurons, but not all of them, innervating each vastus muscle receive common input from a unique source (green and violet lines), but some motor neurons receive inputs from the source directed to the other muscle (dashed green and violet lines; upper graph) and some receive inputs from both sources (lower graph).

The correlation between each motor unit and its neural module reveals the potential nature of shared synaptic inputs by the motor neuron pools that is inevitably obscured in the global EMG signal. With this analysis we show that the motor neurons from two hand muscles during an independent task can by fully retrieved by the module they carry, but the motor units for each knee extensor muscle receives common input from two unique sources.

Previous experiments reported a single dominant common input governing coordination of the vastus medialis and lateralis muscles (26). Similarly, previous studies on one motor unit pool have identified a single common synaptic input (42, 44). The identification of a single dominant common input in previous studies is based on a pooled-coherence approach to estimate neural connectivity. This analysis averages the correlation between motor unit spike trains with several permutations, therefore, the averaging process inevitably generates significant correlations because ~50% of the homonymous motor unit pool receives a similar input. Because we found that the motor units innervating an individual muscle can receive more than one common input, our results demonstrate that pooled coherence is not an appropriate approach to assess neural connectivity.

It is important to note that our experimental task involved isometric contractions in contrast to the dynamic actions typically used to identify ‘muscle synergies’. Perhaps, the sources of common input, such as the type and intensity of feedback from sensory receptors (58, 59), differ during isometric and dynamic contractions and the common input received by the motor neurons innervating the synergistic muscles is more homogeneous. For example, we found that in macaque monkeys during rapid high force contractions most motor units share the same neural module (60).

Even for isometric contractions, however, the sources of common input may differ with the details of the task being performed. Based on the interpretation that the fluctuations in force during steady isometric contractions are attributable to the variance in the common synaptic input (11, 12, 53), differences in the coefficient of variation for force during a specific action suggest adjustments in the common input across tasks. For example, the coefficient of variation for force during index finger abduction, which is mainly due to the activity of the first dorsal interosseus muscle, was 2x greater when participants performed index finger abduction and wrist extension at the same time even though the abduction force was the same in both tasks (61). Based on the finding of an increase in the coefficient of variation for force during the double-action task (index finger abduction + wrist extension) in the study by Almuklass et al. (2016), it seems reasonable to predict that the neural modules for the two hand muscles in the current study would differ from those observed during the independent actions. Consistent with this possibility, Desmedt and Godaux suggested that the synaptic inputs delivered to the motor neurons that innervate the first dorsal interossei muscle differed when the direction of the force applied by the index finger changed from abduction to flexion (62). The basis for this conclusion was the finding that the recruitment order for some pairs of motor units (~8%) consistently reversed recruitment order when the task was changed from abduction to flexion. They hypothesized that this effect, although relatively modest, was attributable to differences in the distribution of the motor command for each task.

Despite the limited scope of the tasks examined in our current study, the findings indicate that the derivation of muscle synergies is based on the common synaptic input that is shared by the motor neurons involved in the action but that this common input is not shared among most of the motor neurons within a given motor nucleus. Moreover, we found that the modulation of discharge rate for all motor units could be classified into three clusters distributed across two neural modules. These results indicate that synergistic motor neuron pools receive common synaptic inputs from at least two different sources during submaximal isometric contractions.

## Methods

### Participants

Eight subjects were recruited for each experiment (hand and knee extensor). All procedures were approved by the local ethical committees at the University of Rome Foro Italico (approval number 44680, knee extension experiments) and Imperial College London (approval number 18IC4685, hand experiments) and conformed to the standards set by the *Declaration of Helsinki*. The subjects signed an informed consent before participating in the study. Some results from these datasets have been published previously (56, 63).

As described subsequently, high-density EMG recordings (Quattrocento, OTBioelettronica, Turin, Italy) and decomposition of the acquired signals (64) were performed in both experiments.

### Experiment 1 (knee extensors)

Participants visited the laboratory on two occasions. In the first visit, they were familiarized with the experimental procedures by performing a series of maximal and submaximal isometric contractions with the knee extensors. In the second visit, which occurred 24 hours after the familiarization session, simultaneous recordings of the force generated by the knee extensors during maximal and submaximal voluntary contractions and HDsEMG signals were recorded from vastus lateralis and vastus medialis. After standardized warm-up contractions, participants were verbally encouraged to push ‘as hard as possible’ for ~3-5 s to achieve peak maximal voluntary force (MVC). They performed ≤4 trials with ~60 s of rest between trials. Approximately 5 min later, they performed steady contractions (2 x 10 % MVC for 70 s) and submaximal trapezoidal contractions at three target forces (2 × 35, 50, 70% MVC force). The trapezoidal contractions required participants to match a prescribed trajectory that comprised a ramp-up phase (5% MVIF s-1), a plateau (10 s of constant force at target), and a ramp-down phase (−5% MVIF s-1). Three minutes of rest was provided between all submaximal contractions. In this study we only used the submaximal steady state contractions at 10% of maximum.

All measurements were performed with both legs with the order determined randomly. Participants were asked to avoid exercise and caffeine intake for 48 hours before testing. Participants were comfortably seated and secured in a Kin-Com dynamometer (KinCom, Denver, CO, USA) by means of three Velcro straps (thigh, chest, pelvis), with the knee joint fixed at 45° of flexion (full knee extension at 0°). HDsEMG signals were acquired from the vastii muscles with two grids of 64 electrodes each (5 columns × 13 rows; gold-coated; 1 mm diameter; 8 mm inter-electrode distance; OT Biolettronica, Turin, Italy) (Fig. 1A). Placement of the electrode grids was based on existing guidelines (Barbero et al. 2012) and adjusted as necessary. After shaving and cleaning the skin (70% ethanol), both electrode grids were attached to muscle surfaces using two layers of disposable double-sided foam (SpesMedica, Battapaglia, Italy). Skin-electrode contact was ensured by filling the holes of the foam layer with conductive paste (SpesMedica). A ground electrode was placed on the contralateral wrist, whereas the reference electrodes for both vastus lateralis and vastus medialis grids were attached to the skin over the ipsilateral patella and medial malleolus, respectively. Monopolar HDsEMG signals were recorded using a multichannel amplifier (EMG-Quattrocento, A/D converted to 16 bits; bandwidth 10-500 Hz; OT Bioelettronica).

### Experiment 2 (hand muscles)

The experimental setup involved a chair, table, and computer monitor. Participants were comfortably seated with both arms resting on the table. A custom-made apparatus that was secured to the table supported the dominant hand (self-reported) in a position midway between pronation and supination and the forearm and wrist were immobilized. The index finger was aligned with the longitudinal axis of the forearm, and the thumb was held in a resting position at the same height as the index finger. The applied force was displayed on a monitor that was positioned 60 cm in front of the subject. The visual gain was fixed at 66 pixels per percentages of MVC force for each muscle (axis). The forces exerted by the index finger and thumb were measured with a three-axis force transducer (Nano25, ATI Industrial Automation, Apex, NC, USA), digitized at 2048 Hz (USB-6225, National Instruments, Austin, TX, USA), and low-pass filtered with a cut-off frequency of 15 Hz. HDsEMG signals were recorded with a multichannel amplifier (OT Bioelettronica Quattrocento, Turin, Italy; bandwidth: 10-500 Hz; resolution: 16 bits) at a sampling rate of 2048 Hz. Two flexible grids of high-density EMG electrodes (13 × 5 pins, 4 mm interelectrode spacing) were placed on the skin over the FDI and thenar muscles (flexor pollicis brevis and abductor pollicis brevis).

Participants performed force-matching tasks (10% MVC force) involving concurrent abduction of the index finger and flexion of the thumb (Fig. 2A). Subjects performed two sustained index finger abduction and thumb flexion contractions for 60 s. Visual feedback was provided as a moving dot cursor in which the x-axis and y-axis corresponded to the thumb and index finger forces, respectively. Subjects had to maintain the force signal within 10% of the target.

The experiments began with MVCs (as described in Experiment 1). After the MVCs were determined, the required target forces were displayed on a monitor. Participants performed two 60-s trials with a 30 s of rest between trials. As noted in the introduction, we designed our tasks to determine the extent to which distinct motor neuron pools would receive common inputs. To achieve this goal, subjects were instructed to exert forces in the same sagittal plane, which required ~10 minutes of practice.

### Data analysis

The 64 monopolar HDsEMG signals were filtered offline with a zero-lag, high-pass (10 Hz) and low-pass filter (500 Hz). The force signals were corrected for the influence of gravity and normalized to MVC force. HDsEMG channels with poor signal-to-noise ratios were inspected with a semi-automated function that identified spurious EMG signals based on the power spectrum. Those channels with a poor signal-to-noise ratio (defined as 3 standard deviations from the mean, power spectrum averaged across all signals in the band 10 - 500 Hz) were visually inspected and removed from the analysis. The number of EMG channels containing noise was low; > 95% of the channels had good signal-to-noise ratios.

Subsequently, the HDsEMG signals were decomposed with a gradient convolution kernel compensation algorithm (48). The general decomposition procedures have been described previously (49). Briefly, the EMG signals can be described as time-series of Dirac delta functions that contain the sources (s) representing the discharge times of motor units. The time series of the motor unit discharge times can be described as delta (*δ*) functions:

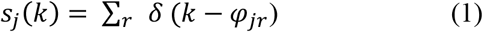

where *φ_jr_* corresponds to the spike times of the j*th* motor unit. Each channel of the EMG signal can be then described as convolution of the motor unit discharge times (s) into the muscle fiber action potentials. Because each motor unit innervates multiple muscle fibres, it is possible to observe a compound action potential from the muscle fibers innervated by that motor axon. Therefore, the HDsEMG recordings can be described mathematically in a matrix 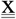 form as:

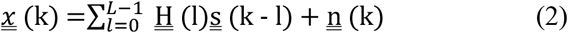

where 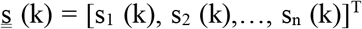 represents the n motor unit discharge times derived from the EMG signal 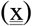 and 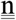 is the noise to for each electrode. The matrix 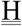 contains the two-dimensional information of the motor unit action potential and has size m x l with l*th* sample of motor unit action potentials for the *n* motor units and *m* channels.

Before the beginning the blind source separation procedure, the spatial sparsity of the matrix x was enhanced by extending the observation numbers. This procedure improves the decomposition as the gradient descent update rule maximises the diversity of the motor unit waveform to converge the discharge times of each motor unit (the sources, s). Because this process is blind, it is possible to inspect the shapes of the motor unit action potentials obtained by spike triggered averaging and to perform visual inspections of the 2D and 3D waveforms (see 50, 60).

### Factorization analysis

The neural control of muscles by motor neurons can be described and predicted analytically. If the discharge times of the motor units are known, it is possible to predict modulation of muscle force with near perfect correlations (52, 65). By recording of large samples of motor units, it is possible to reconstruct modulation of muscle force (11) due the low-pass filtering properties of the muscle to a given neural drive (51, 52). When motor unit discharge rates are filtered in the low-frequency range (muscle bandwidth <20 Hz), it is possible to predict oscillations in the applied force close to ~1% MVC (51). Consequently, the factorization analysis used in the current study focussed on the low-pass filtered motor unit discharge rates. The discharge rates were filtered by convolving with a Hann window of 400 ms (2.5 Hz). The motor unit discharge times were factorized with three methods: factorization analysis (66, 67), principal component analysis, and non-negative matrix factorization (see Figure 1 in supplementary materials).

These factorization methods were applied on the matrix containing the smoothed discharge rates with rows equal to the number of motor units identified for both muscles and columns equal to the smoothed discharge times. Figures 1 and 2 shows examples of this procedure for the vastii and hand muscles.

Factorization analysis is based on the rationale that muscle force is the consequence of the activation of many motor units, which can be represented as time sequences of M dimensional vectors (see equation 1) due to the activation of the motor neurons **m(t)** in response to various common and independent synaptic inputs. Thus, the response of the motor neuron population can be described as combinations of N varying synaptic inputs that are constrained by the non-linear properties of the motor neuron, which construct a specific motor unit characteristic, or *neural module*, expressed as ({*w_i_*(*t*)}_*i*=1,…*N*_

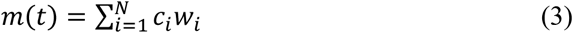

where *c_i_*, is a non-negative scaling coefficient of the *i*-th neural module. We were interested in finding the matrix *w_i_* without making any assumptions about the relations between muscles or motor neurons. We found that factor analysis was the best method in terms of correlations of the neural modules to the discharge times of individual motor units. Moreover, we demonstrate with an integrate-and-fire model (see below) that factor analysis can separate the neural modules with high levels of accuracy. We also examined the performance of non-negative matrix factorization and principal component analysis by using previous approaches to identify muscle synergies (i.e., >100 iterations and reconstruction of the original signal, (19, 20, 25)).

The factor analysis models the associations between variables into fewer latent variables (factors). It assumes that for a collection of observed variables (*x*) there are a set factors (*f*) that explain most of the total variance, which is the common variance. The function *factoran* (in Matlab) computes the maximum likelihood estimate of the factor loadings matrix (Λ)

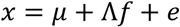

where *e* is the vector of independent specific factors. Alternatively, the model can be specified as

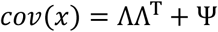

Where ΛΛ^T^ is the common variance matrix and Ψ = *cov(e)* is the diagonal matrix of specific variances. The unique variance in the model with no a priori assumption of orthogonality between factors (when allowing for factor rotations such as *promax*) makes the factor analysis an appropriate choice to extract the latent discharge rate of the synergistic motor nucleus. It is supposed that the model mimics the common and independent inputs impinging into the two motor nuclei.

### Crosstalk and realigning

Motor unit action potentials from the first dorsal interosseous into the thenar muscles (and vice versa) and from the vastus medialis into vastus lateralis can experience cross-talk up to 95%; that is, 95% of motor units from one muscle being conducted with minimal shape distortion to the neighbouring muscle (55). Consequently, we examined the level of cross-talk with a validated method (55, 56). Briefly, this procedure takes advantage of the distance from the activated muscle fibers (muscle unit) to the electrode, which is less for motor units in a targeted muscle. Motor units from a targeted muscle are expected to show greater action potential amplitudes with minimal shape distortion (action potential conduction velocity in the grid, see Germer et al. 2021 for the detailed assessment of statistically significant crosstalk of individual motor units).

Another step was to realign motor unit action potentials with respect to the averaged motor unit action potential that was obtained after spike-triggered averaging. Because the action potential at individual time instants shows some variability due to surface EMG stochasticity, we convoluted the average action potential to retrieve the time instants of activation of the motor units. Although this procedure was critical for assessing accurate brain-spinal transmission latencies (68), it did not influence our results because corticospinal latencies are so small (<50 ms). In our study, the firing rate was smoothed in the frequency of force production (2.5 Hz, 400 ms). The effects of action potential onset timing, therefore, are removed when low-pass filtering the motor unit discharge timings at <2.5 Hz.

### Computer simulations

We simulated 480 integrate-and-fire motor neurons, each of which received a computer-generated current input (set at 20 nA). The synaptic currents that were shared among all neurons to represent the common synaptic input, but the neurons also received some independent synaptic inputs. Because motor neurons can exhibit synchronous discharge times, the common input currents were close to maximal values in the low frequency range <2.5 Hz (see below). The resting membrane potential and reset voltage was set at −70 mV, the spike threshold was set at −50 mV, and a membrane time constant was 20 ms. The timestep duration was set at 0.1 ms.

Our model comprised randomly distributed gaussian noises at each time step to represent the common and independent synaptic inputs. Figure 6 shows the overall architecture of the model. Two random uncorrelated gaussian input currents were created at each time step to represent the *neural modules* that were identified experimentally. One-third of the neurons (160 neurons) received *I*_syn1_ as a unique common input. *I*_syn2_ received the same common input strength but orthogonal to *I*_syn1_. *I*_syn3_ was the average of *I*_syn1_ and *I*_syn2_ plus its unique independent noise (see eq. 5). The input current for each neuron *i* and population *j* (*j* = 1,2,3) can be summarized by the following equation:

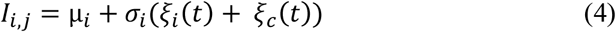

where μ_i_ is the temporal average of the current that was set at 20 nA and σ_i_ sets the global network state by taking into account the unique independent inputs for each cell (*ξ_i_*) and the gaussian-distributed random common inputs (*ξ_c_*) The tuning of these parameters was matched to those observed in vivo. The values of μ, *ξ_i_*, and *ξ_c_* were adjusted to reflect physiological values for the variability in motor unit interspike intervals and common input. Interspike interval variability was examined with histogram distributions, as found in the current study and by others (69). The common and independent inputs were tuned based on the cross-correlogram function derived from previously reported motor unit data (44). Therefore, each combination of three randomly assigned groups of 160 neurons from the total pool (n=480) received two independent synaptic currents (*I*_*i*1,2_ equal to equation 4) and a third randomized subpopulation (*j* = 3) of motor neuron received the average of the two inputs:

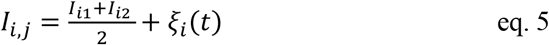

The bivariate correlations between the synaptic currents are shown in Figure 5. The total duration of the simulation was set at 10, 50, and 80 s. We removed the first and last 2 s of spiking activity for all simulations to minimize the influence of spike-frequency adaptation. The spike trains emitted by the *Ith* neuron after generating the spike times were stored as a binary time series, which was equal to 1 when the neuron reached voltage threshold. The successive analysis followed the same steps as the experimental data. Briefly, the binary spike trains were low-pass filtered with a 2.5 Hann window and the factorization analysis was then performed on the low-pass filtered signals. Because the distribution of inputs to each neuron were known (*i.e*., *I*_*i*1-3_), it was possible to retrieve the performance accuracy of the factorization analysis. Moreover, it was possible to investigate the relation between motor neuron responses to increased synaptic currents with changes in common and independent inputs. As reported in the results, we found that when a large number of spikes was included in the analysis (simulation of 80 s), the factorization analysis provided a perfect prediction of the three sources of common synaptic inputs. Our model confirmed that the three clusters observed experimentally in the neural modules arise due from two distinct oscillatory common synaptic inputs and that the third component (the shared neural modules) is an average of the two other modules.

### Statistical analysis

We performed linear regression analysis between the smoothed motor unit discharge rates within and between muscles. The significant level was extracted from bivariate Pearson’s correlations and Bonferroni method was applied with multiple comparisons. The same procedure was used to find the modules carried by each neuron (decoding function). Each neural module extracted by the factorization algorithm (66, 67) was compared to the firing rate of the individual motor units. Significance was accepted for P values < 0.05.

## Supplementary materials

**Figure 1S.**
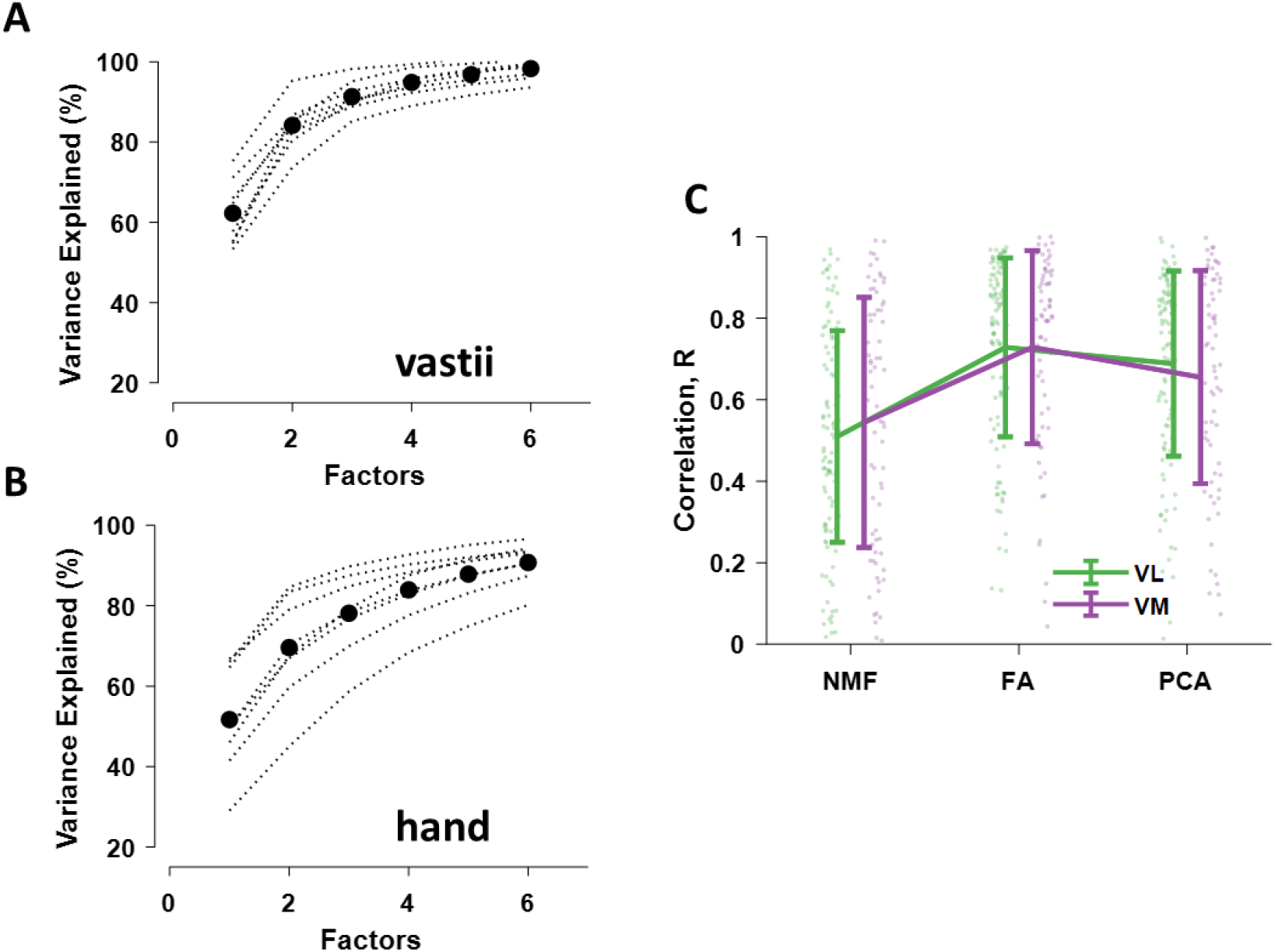
Reconstruction accuracy (% variance explained) for each subject (dotted black lines). The black dots in **A** and **B** represent the average neural modules across subjects. **C**. The correlation values (mean ± SD) between the modules and the motor units discharge rates.

